# Tinamou egg color displacement at eco-geographical and song space overlap

**DOI:** 10.1101/2022.09.27.509793

**Authors:** Qin Li, Dahong Chen, Silu Wang

**Affiliations:** Department of Science and Education & Grainger Bioinformatics Center, Field Museum of Natural History, Chicago, IL 60605, USA; Nuclear Organization and Gene Expression Section, Laboratory of Biochemistry and Genetics, National Institute of Diabetes and Digestive and Kidney Diseases, National Institutes of Health, 9000 Rockville Pike, Bethesda, MD 20892, USA; Department of Evolution and Ecology, University of California, Davis, 95618

## Abstract

The divergence of reproductive traits frequently underpins the evolution of reproductive isolation. Here we investigated the hypothesis that tinamou (Tinamidae) egg coloration functions as a mating signal and its diversification was driven by reinforcement. For many tinamou species, the male guards the nest that is sequentially visited and laid eggs in by multiple females. The colorations of the existing eggs in the nest could signal mate quality and species identities to both the incubating male and the upcoming females, preventing costly hybridization, thus were selected to diverge among species (Mating Signal Character Displacement Hypothesis). If so, two predictions should follow: (1) egg colors should coevolve with known mating signals as the tinamou lineages diverged; (2) species that partition similar eco-geography should display different egg colors. The tinamou songs are important mating signals and are highly divergent among species. We found that the egg color was significantly associated with the first principal component of the song variables. In addition, tinamou species with similar eco-geography tend to display different egg colors, while controlling for song variation among species. Mating signal evolution could be opportunistic and even exploit post-mating trait as premating signals that undergo character displacement in sympatry.

## Introduction

Reproductive trait divergence is crucial for speciation because these traits frequently underpin barriers to gene flow in the onset of speciation (Koski and Ashman 2016; Pfennig 2016). A very puzzling reproductive trait divergence exists in tinamou egg coloration. Tinamiformes (common name: tinamou) is the most species-rich order of Palaeognathae, containing 48 extant species (Figure 1) (Cabot 1992; Davies 2002). Tinamous are mostly dull in plumage, but various species of tinamous lay brightly colored eggs ranging from brilliant magenta, to pink, purple, turquoise, and olive green (Figure 1) (Cabot 1992). Despite its evolutionary uniqueness, tinamou egg color divergence remains a mystery due to the cryptic and elusive plumage and behavior of the birds (Cabot 1992; Davies 2002).

**Figure 1.**
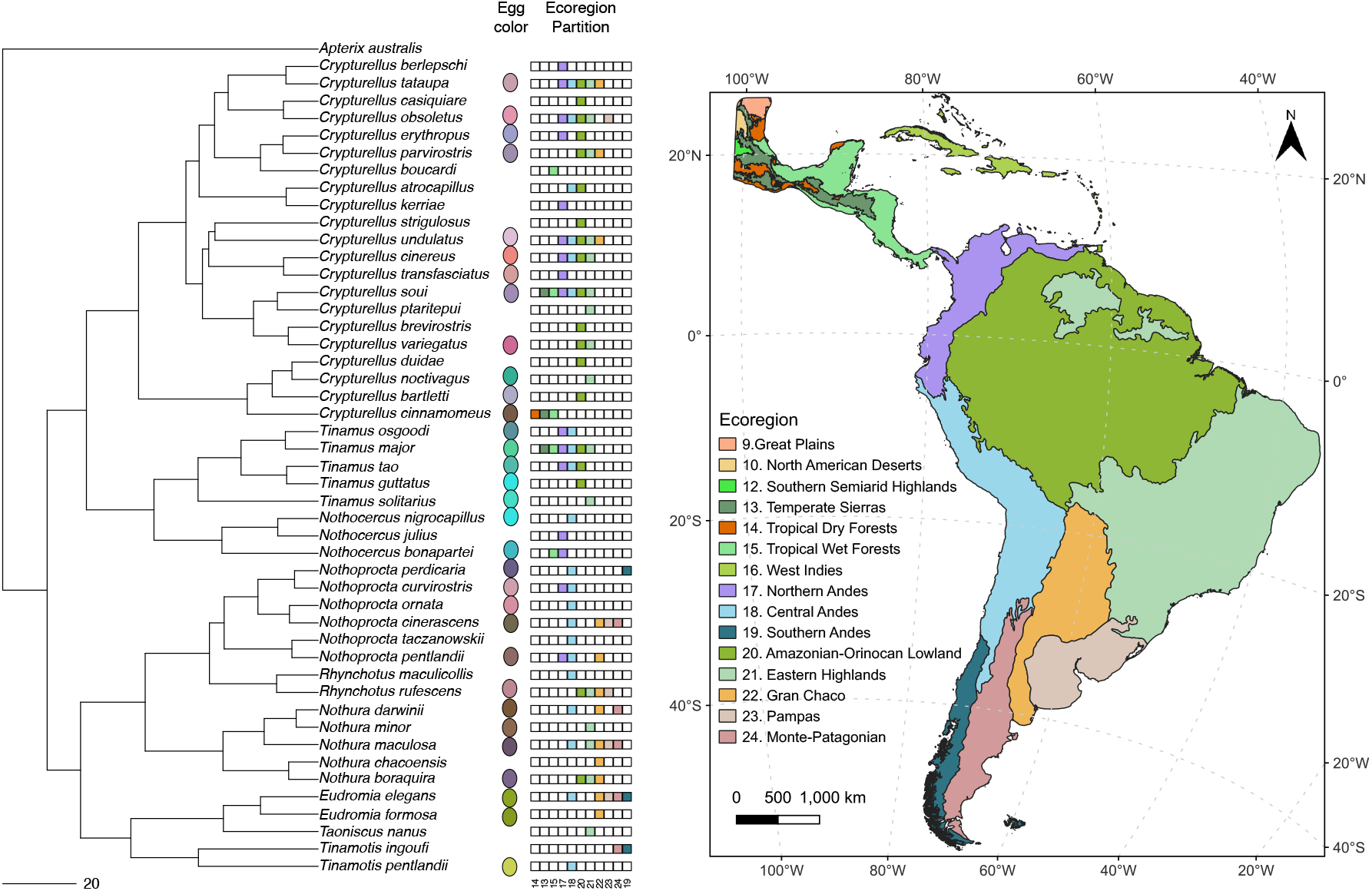
Egg color and ecoregion partition across the Tinamidae phylogeny. Tinamou ecoregion and song space co-partitioning is significantly associated with egg color difference among species, with greater color divergence among species that partition similar ecoregions after controlling for song variation (partial mantel test, *p* < 0.05). The Tinamidae phylogeny (with *Apterix australis* as the outgroup species) was inferred from OneZoom Explorer (http://www.onezoom.org). The Lambert azimuthal equal-area projection was applied for ecoregion map.

Two main hypotheses have been made about evolutionary mechanism of egg color divergence: (1) ‘aposematism hypothesis’ (Swynnerton 1916) states that eggs are brightly colored to warn predators of their distastefulness; (2) ‘mating signal hypothesis’, which predicts that egg color are employed for stimulating reproductive investment of the incubating sex (Weeks 1973; Brennan 2010). However, ‘mating signal hypothesis’ (2) did not explain egg color divergence among tinamou species. We further generalize the ‘mating signal hypothesis’ (2) into “mating signal character displacement hypothesis’ (3), in which the egg colors could signal to both sexes for mate recognition and attraction, that could be divergently selected among closely related species in sympatry. The aposematic function of egg coloration was thought to be unlikely, because egg predators are nocturnal mammals or reptiles that prioritize chemical cues over visual cues (Skutch 1966; Cabot 1992; Brennan 2010). The ‘mating signal character displacement hypothesis’ (3) is plausible considering the ecology and mating systems of tinamou species. Although there is limited understanding of tinamou mating systems, for the species of which mating systems were described, males collect and incubate eggs laid by multiple females (Cabot 1992; Davies 2002). An initial female is attracted by distinctive songs to the nest guarded a male. If mating occurs, the female lays eggs in the nest and leaves before other females come and lay more eggs to be incubated by the same male (Cabot 1992; Davies 2002). The colors of the existing eggs in the nest could signal to both the incubating male and the upcoming females. As predicted by the mating signal hypothesis (2), the salient egg colors could stimulate male incubation and reproductive investment, as well as convey intra-specific mate identity. We further suspect that the egg colors could signal to the upcoming females to the focal nest as mate-copying and species-recognition cues. Such alternative “mating signal channel” might be especially favored when the plumage is adapted for camouflage (Cabot 1992; Davies 2002). If egg colors of different tinamou species is adapted for mate recognition, reinforcement (Dobzhansky 1937; Mayr 1942; Liou and Price 1994; Servedio 2000) could have driven egg color divergence among different tinamou species.

Here we investigated the ‘mate-signal character displacement hypothesis’ (3) of tinamou egg color divergence. If tinamou egg colors are employed as a species-specific mate recognition signal, it should coevolve with other mate-recognition signals as the tinamou lineages diverged. Songs of tinamous are known to be important and highly divergent mating signals among species (Cabot 1992; Bertelli and Tubaro 2002; Laverde-R. and Cadena 2014; Boesman et al. 2018). In addition to songs, multimodality of mating signals is selected for efficient mate-recognition in complex environment in a number of taxa (Partan and Marler 1999; Rowe 1999; Secondi et al. 2015; Halfwerk et al. 2019). When glamorous plumage is costly due to increased predation risk, alternative signal modality such as egg coloration could be favored. Since the birds themselves (instead of the eggs) tend to attract nest predation (Brennan 2010), egg coloration could be employed to fulfill signal multimodality, bypassing the evolutionary constraints for plumage. If the egg colors provide an additional dimension of mating signal, tinamou species with similar songs and ecogeographical range should have different egg colors. To investigate this possibility, here we ask: (1) whether egg coloration coevolved with songs as tinamou species diverged; and (2) whether species that are more likely to have a sympatric history and similar songs display more divergent egg colors?

## Methods

To investigate the mating signal character displacement hypothesis in tinamou egg color evolution, we examined the association of song, egg colors, and ecoregion co-partitions among tinamou species.

### Tinamou egg color

Tinamou egg coloration data was extracted from existing tinamou egg color analysis in an online article (Schläpfer 2017), which quantified coloration of the eggs in RGB color space from 32 tinamou species based on nest photo citizen science submissions. Since tinamou egg colors are known to decay over the course museum storage, tinamou nest photos are used to best represent the functional egg color among species (Schläpfer 2017). Briefly, for each species, the egg coloration reflected in RGB color space was quantified (Schläpfer 2017). To efficiently infer egg color variation among species, we conducted principal component analysis (PCA) of the RGB color space with RGB axes (values centered around zero and scaled to unit variance).

Ideally, egg colors should be quantified in the nest in the tetrahedron color space that is consistent with tinamou vision. However, given the difficulty in nest detection in the clade of elusive avian species, Schläpfer (2017) resorted citzen science nest photos and quantified species-specific egg colors in RBG color space. A caveat of this metric is the lack of detection in color variation in the ultraviolet (UV) axis and noneuqal-distance conversion to the tetrahedron space. However, this might not be an problem for tinamou visual system, as an earlier study suggested that tinamous likely do not have UV perception since neither the brushland tinamou (*Nothoprocta cinerascens*) nor the Chilean tinamou (*Nothoprocta perdicaria*) had short wavelength sensitive type1 cones (Mullen and Pohland 2008). Further, RGB color metric has been shown to be highly predictive to avian perception (R^2^ = 0.837) (Bergeron and Fuller 2018) and thus provides an effective and practical way to detect egg color variation among tinamou species given the challenges in egg-searching, curation, and color preservation.

To test the validity of between-species variation in the egg color quantification in RGB space from field nest photos (Schläpfer 2017), we compared the within-versus between-species egg color differences with museum egg specimens of four tinamou species: *Crypturellus undulatus, Eudromia elegans, Nothura boraquira, Tinamus major*. Photos of egg collections from the Field Museum of Natural History were taken with a standard color reference chart (by Canon EOS 5D Mark IV; Figure S1 B). All images were set to the RGB 16bit mode in Photoshop. All images were white-normalized to the first white patch of the standard. Central areas 3mm away from the edge of each egg were selected, and all reflection, cracked slots and labels were excluded from further quantification. Colors of the remaining central areas were averaged to generate one RGB reading for each egg. Up to four eggs at the most left and top positions were quantified if more than four eggs from a clutch were photographed in one image. Accounting for the egg color variation over time within species, we tested whether there are species-specific egg color clusters with both field nest photo (Schläpfer 2017) and our museum photo measurements for all the four species.

### Tinamou song

Song data was acquired from a previous study (Bertelli and Tubaro 2002), in which four song variables from 40 tinamou species were quantified: maximum frequency (Hz), minimum frequency (Hz), emphasized frequency (frequency of the note with highest amplitude in the song, Hz), and bandwidth (the difference between maximum and minimum frequency, Hz). Because these variables are correlated (Figure S2), we used PCA with scaled values to generate PC1 that captured 83% of the variation in song data. Since previous study showed that song covaries with habitat (Bertelli and Tubaro 2002), we have to consider habitat variation in the current study.

### Tinamou ecoregion-partitioning

Species ranges were generated according to their occurrences. Geo-referenced occurrences were downloaded from GBIF (GBIF.org (30 September 2021) GBIF Occurrence Download https://doi.org/10.15468/dl.fqnhrk) for all tinamou species. Occurrences with uncertainty in coordinates less than 10km were kept, and were further filtered by removing redundant localities within a 1-km^2^ grid cell via R package *ecospat* (Cola et al. 2017) after Lambert azimuthal equal-area projection (ranging 9 – 9413, median = 469). Ecoregion designation for each species was carried out by overlaying occurrences with the ecoregion map of America. Ecoregion maps were downloaded for North America (https://www.epa.gov) and South America (http://ecologicalregions.info) separately at level I and then were combined as one to include 11 ecoregions for all occurrences. For each species, an ecoregion within which the proportion of occurrences was greater than 10% (or > 50 if the total number > 1000) was included in the species’ ecoregion distribution.

With this data, we calculated pairwise probability of ecoregion co-partitioning among tinamou species. For each pair of species, the probability of ecoregion co-partitioning is the sum (across all ecoregions) of the multiplication of the probability of species-specific partitioning within each ecoregion. For each species, the probability of partitioning within an ecoregion is the number of points of occurrence within the ecoregion over the total points of occurrence for the species. If the probability of ecoregion co-partitioning is an effective indicator of sympatry or parapatry in the course of tinamou speciation, closely related species should be more likely to co-partition ecoregions, given the limited dispersibility of tinamous (Davies 2002). We employed Mantel Pearson’s correlation test with function *mantel*.*test* in R to test if the species distance matrix is correlated with the ecoregion co-partitioning probability matrix.

### Tinamou phylogeny

To account for phylogenetic uncertainty, we downloaded a phylogeny subset of 500 random trees with Hackett backbone for tinamou species from VertLife (http://vertlife.org/phylosubsets; Jetz et al. 2012). Additionally, we generated a representative phylogeny based on OneZoom Explorer with branch length as divergence time for visualization (http://www.onezoom.org).

### Statistical tests

We tested four macroevolutionary associations: (1) egg color and song; (2) song and habitat; (3) egg color and habitat; and (4) egg color displacement at ecoregion and song space (song PC space) co-partitioning among tinamou species. Species-specific habitat type is a variable with “open”, “mixed”, and “close forest” habitat categories acquired from previous study (Bertelli and Tubaro 2002). For (1)-(3), we used *gls* with *corPagel* function in the *ape* package R to account for phylogenetic signals (Paradis and Schliep 2019). For egg color and song variables, we used PC axes that cumulatively explain over 80% of the variation, which resulted in PC1 (54%) and PC2 (38%) for egg color and only PC1 (83%) of song PCA. To account for phylogenetic uncertainty, we used 500 tinamou phylogenies. To correct for multiple hypotheses testing, we conducted False Discovery Rate correction (Benjamini and Hochberg 1995).

Further, we examined whether tinamou egg colors are character displacement trait, by testing whether there are greater egg color differences between species that are more likely to co-partition eco-regions. Since song can be a character displacement trait as well, we have to control for song disparity among species. We first computed the distance matrices of egg color PC1 and song PC1 respectively with the *dist* function in R. To examine the correlation between the distance matrix of egg color PC1 and the probability matrix of ecoregion co-partitioning while controlling for song PC1 distance matrix among species, we used one-tailed Partial Mantel Pearson’s correlation test with 10,000 iterations with the *mantel*.*partial* function. All the variables involved in the study is deposited on Dryad (https://doi.org/10.5061/dryad.rfj6q577g).

## Results

Egg color is significantly associated with song and habitat variation among tinamou species (Figure 2 A). The between-species distinction overcomes the within-species variation in egg color variation (Figure S1). Ecogeographic similarity predicts egg color distance among species, after controlling for song variation among tinamou species (Figure 1).

**Figure 2.**
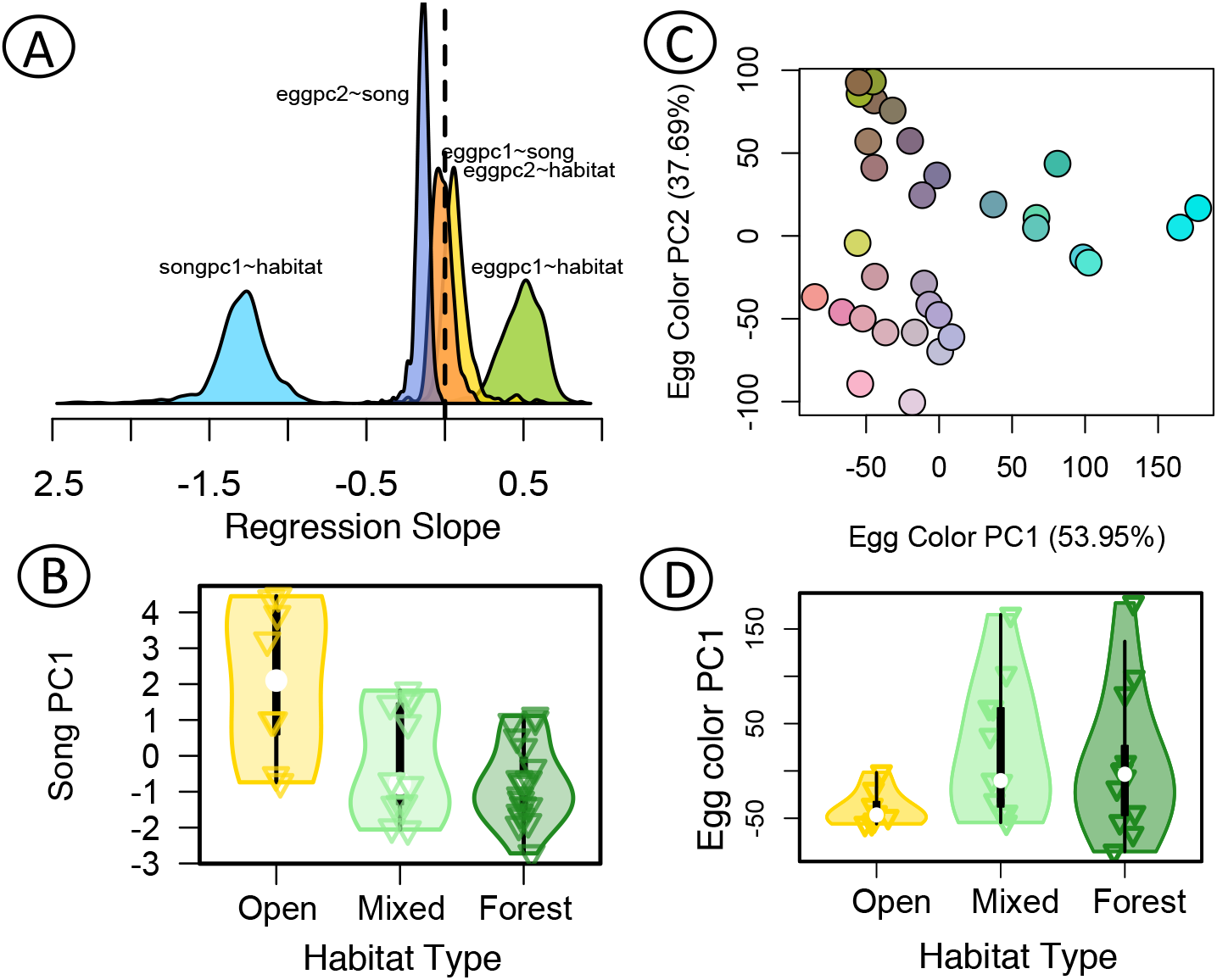
Association of habitat type, song variation, and egg color among tinamou species. (**A**) The distribution of regression slope between response variables and predicting variables derived from 500 tinamou phylogenies (Table 1). Three distributions of regression slope were significantly deviated from zero: (1) between song PC1 and habitat type (**B**), (2) between egg color PC1 (**C**) and habitat (**D**), as well as (3) between egg color PC2 and song PC1 (**A**). (**B**) Phylogenetic analysis reveals significant association between song PC1 and habitat type among tinamou species, consistent with a previous study (Bertelli and Tubaro 2002). (**C**) Egg color PC1 and PC2: the dot color represents the egg RGB color of each species. (**D**) Egg color PC1 is significantly associated with habitat types among tinamou species.

Egg color is associated with song and habitat variation specifically (Figure 2). Among 32 tinamou species, PC1 reflects the warm-to-cold gradient of egg colors, while PC1 reflects the brightness (Figure 2 C). There was significant association between egg color PC2 and song PC1 (Table 1; Figure 2 A). The relationship is unlikely confounded by habitat types among tinamou species, as egg color PC1, but not egg color PC2, was significantly associated with habitat types (Table 2; Figure 2 C-D). The open field tinamou species tend to lay warm-colored eggs, while there was a wider variation in egg colors (from pink to greenish blue) among the closed forest tinamou species (Figure 2 D).

**Table 1.** Phylogenetic regression models investigating the associations among egg color, song, and habitat types across tinamou species. For each pair of variables, a phylogenetic regression model was run with each of the 500 tinamou phylogenies to account for phylogenetic uncertainty. The significant (0 falls out of the 95% confidence interval, CI) intercepts (*b*_*0*_) and slopes (*b*_*1*_) were in bold.

| Response variable | Predictor | mean $b_1$ | 95% CI $b_1$ | mean $b_0$ | 95% CI $b_0$ | R<sup>2</sup> |
| --- | --- | --- | --- | --- | --- | --- |
| egg color PC1 | song PC1 | -0.017 | (-0.118, 0.128) | -0.007 | (-0.274, 0.265) | 0.008 |
|  | habitat | **0.483** | (0.151, 0.693) | **-1.074** | (-1.546, -0.353) | **0.119** |
| egg color PC2 | song PC1 | **-0.144** | (-0.244, -0.072) | **0.157** | (0.034, 0.278) | **0.069** |
|  | habitat | 0.071 | (-0.090, 0.392) | -0.060 | (-0.750, 0.259) | 0.010 |
| song PC1 | habitat | **-1.297** | (-1.729, -0.990) | **2.929** | (2.175, 3.861) | **0.383** |

Ecoregion co-occurrence is likely an effective indicator of parapatry or sympatry in history of tinamou speciation, as closely related species tends to co-occur at ecoregions (Mantel Z = 1530.38, *p* = 0.001). We further observed greater egg color divergence between species that co-partition ecoregions after controlling for song variation among tinamou species. The PC1 explains 54% variation of egg colors in the RGB color space. The distance of egg color PC1 is significantly positively associated with the ecoregion co-partitioning among species controlling for song distance among tinamou species (Partial Mantel r = 0.066, *p* = 0.048).

## Discussion

The macroevolution relationship among egg color disparity, song variation, and habitat and ecoregion partitioning, is consistent with the ‘Mating Signal Character Displacement Hypothesis’ that tinamou egg coloration is an alternative mating signal that is divergently selected among species with similar ecogeographical partition (Weeks 1973; Brennan 2010). Egg colors and songs could be divergently selected as multimodal mate-recognition signals among tinamou species with similar appearance that partition similar ecogeographical range. When the plumage modality is constrained by anti-predation adaptation, egg coloration could be opportunistically adopted to fulfill mating signal multimodality for species recognition in sympatry. This study sheds light on the evolution of multimodal sexual signals that bypasses natural selection for plumage camouflage in the most species-rich order of Palaeognathae.

Egg coloration could be both a pre- and post-mating signals in tinamous. In many other bird species, egg color is known to be post-mating signals indicating female quality and can influence male incubation and promiscuity (Soler et al. 2005; English 2009). This is consistent with the Mating Signal Hypothesis in which egg colors could stimulate tinamou male incubation and parental care (Weeks 1973; Brennan 2010). In addition, egg coloration in the nest can be premating signals received by upcoming females. Premating mate-recognition signals among females that mate with the same male could be selected if it enhances the offspring fitness by protecting the reproductive investment of the parental care provider (male tinamous) from costly heterospecific mating interference (Gröning and Hochkirch 2008).

Although the mating systems are not well-understood for elusive tinamous, in the species with mating system records, males guard and incubate the eggs laid by multiple females (Cabot 1992). The coloration of the existing eggs in the nest could be a mating signal received by upcoming females. As premating signals, egg colors could reflect mate species-identity and quality (Sætre et al. 1997; Servedio and Noor 2003; Secondi et al. 2015), as well as stimulate female mate choice copying (Dugatkin 1992; Gibson and Höglund 1992).

Then why do tinamous adopt egg colors as mating signals in addition to songs? Songs that are important mating signals in tinamous (Bertelli and Tubaro 2002) and are involved in duetting between mating partners (Boesman et al. 2018). However, such acoustic modality is usually insufficient for mate communication in complex environments, thus signal redundancy or multimodality is further selected for mate communication to alleviate mate-searching effort and/or hybridization (Partan and Marler 1999; Rowe 1999; Uy et al. 2008; Secondi et al. 2015). Plumage patterning and coloration are frequently adopted as inter-specific and intra-specific mating signals at a finer spatial scale in birds (Uy et al. 2008, 2009; Seddon et al. 2013). However, most tinamou species exhibited cryptic plumage as an adaption for anti-predatory camouflage (Cabot 1992; Davies 2002). Sexual selection for mating signal elaboration would compromise the adaptation for camouflage favored by natural selection (Heinen-Kay et al. 2015). Egg coloration can be an alternative modality of mate signaling enrichment towards mate-searching refinement, bypassing the conflict between natural and sexual selection. This idea is supported by the fact that egg color brightness is correlated with song variation among species (Table 1, Figure 2 A-B).

Reinforcement may have driven egg color and song divergence among sympatric or parapatric tinamou species. Many closely-related tinamou species demonstrate historical and/or contemporary ecoregion overlap (Cabot 1992) (Figure 1). Ecological and/or intrinsic incompatibility among diverged lineages leads to reduced hybrid fitness, which in turn select for premating signal divergence to avoid costly hybridization (Dobzhansky 1940; Mayr 1942; Liou and Price 1994; Servedio 2000). Mating signal multimodality is needed to ensure mate recognition in complex heterospecific environment (Uy et al. 2008; Secondi et al. 2015). The divergence of tinamou song and egg coloration could jointly reduce heterospecific reproductive efforts among sympatric tinamou species. Closely related tinamou species that are mostly allopatric at present but could still harbor footprints of historical character displacement of egg colors formed at historical sympatry either over secondary contact or sympatry speciation.

Notably, the species with greater likelihood of ecoregion co-occurrence tend to display divergent egg colors (Figure 1). For example, the closely related species *Crypturellus noctivagus, C. bartletti, C. variegatus* still partition similar ecoregions and demonstrate distinct egg coloration, which may have been driven by reinforcement at historical sympatry (Figure 3). The association of egg color and habitat types among tinamou species (Table 1, Figure 2 AD) further indicates residual divergent selection on egg colors in the speciation history. Although the speciation event among tinamou species have long passed (> 7 million years), the likelihood of ecoregion co-occurrence is potentially an effective indicator of historical sympatry/parapatry in the history of tinamou speciation because the closely related species tend to share ecoregions.

**Figure 3.**
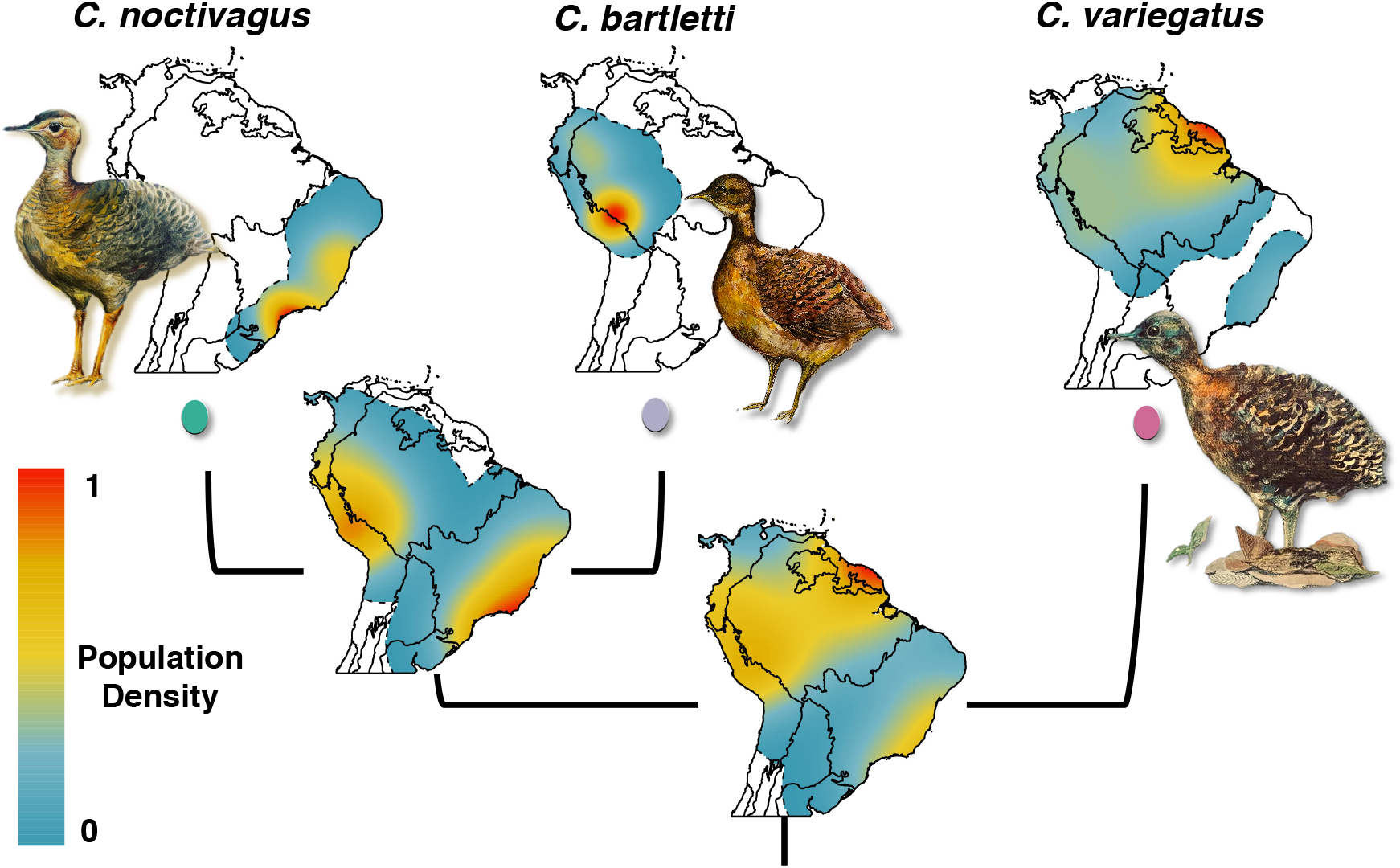
Reconstructed ancestral distribution of three *Crypturellus* species. Black lines depict the boundaries of ecoregions corresponding to Figure 1, and dashed lines depicted boundaries of species’ distribution ranges. Ranges of tips were concave hull polygons generated via R package *rangemap* (Cobos et al. 2021) with a hierarchical clustering method, and kernel densities were estimated via R package *spatstat* (Adrian Baddeley, Ege Rubak 2015), to capture the spatial configuration of the occurrences. Distributions of internal nodes were generated by the same method with an intermediate step of rescaling densities (at a 100-km-resolution) of all descendants to offset the effect of variation in abundance. Blue to red colors depict the relative scale of kernel densities from low to high (ranging 0 to 1).

Although we found significant supports for both predictions of the Mating Signal Character Displacement Hypothesis, the significance levels were marginal. This indicates that there are other potential evolutionary forces shaping tinamou egg color divergence. Besides the local tinamou species assembly, other complex aspects of abiotic and biotic interactions such as sexual conflicts, predation, and parasitism, might contribute to the egg color signal diversity.

The genetic underpinning of such song and egg color association is unclear. The simplest genetic mechanism underlying such association is pleiotropy (one gene affecting multiple traits) (Fisher 1930; Williams 1957). For example, the pleiotropic *foraging* gene underpins the association among social behavior and life history traits in natural populations (de Belle et al. 1989; Mery et al. 2007; Wang and Sokolowski 2017). There might be an omnipotent pleiotropic gene linking egg coloration and songs, among many other traits. If so, a wide array of traits underpinned by such pleiotropy are expected to coevolve. However, the association between song and egg color of tinamous is specific: only brightness egg color is associated with tinamou songs (Table 1, Figure 2 A-B). Such specificity in song-egg-color association is consistent with mating signal multimodality in which specifically signals were coupled among modalities (Gilliard 1956; Partan and Marler 1999; Hebets and Papaj 2005). Various signal modality can be underpinned by different genes and selected to coevolve as multimodal signals. Future investigation of genetic underpinnings of tinamou song and egg colors would further shed light on this interesting co-divergence. In sum, the results presented herein are concordant with the hypothesis that egg coloration in tinamous serves as mating signals, for mate attraction and/or mate species-recognition and could be divergently selected upon historical sympatry/parapatry.

## Acknowledgement

Thank John Bates and Chad Eliason for accessing tinamou egg collections and photographing at the Field Museum of Natural History. Thank Patricia Brennan, Daniel Hanley, Julia Clarke, Jonathan Rolland, Darren Irwin, Graham Coop, and the Coop Lab for helpful discussions. Thank Daniel Hanley, Ken Thompson, Darren Irwin, and Jonathan Rolland for comments on the manuscript.

## Supplementary information

**Figure S1.**
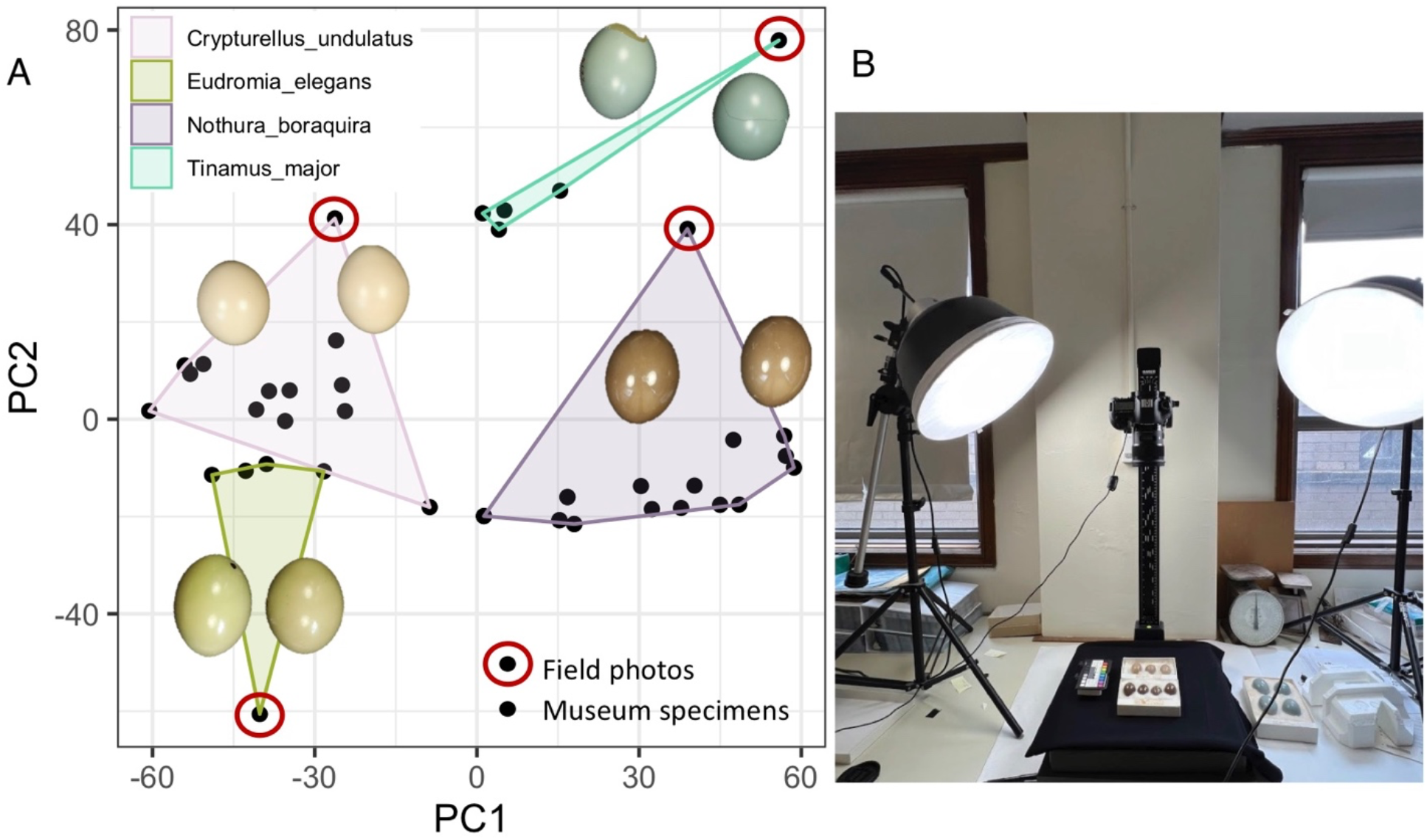
Fisher Discriminant Analysis of egg color. **A**, Egg colors were quantified in RGB space from field nest photos and museum collection within and between tinamou species. Although there is variation within species and between field and museum RGB quantification, the between-species difference is distinctive. **B**, Lighting setup at which the museum egg photos were taken.

**Figure S2.**
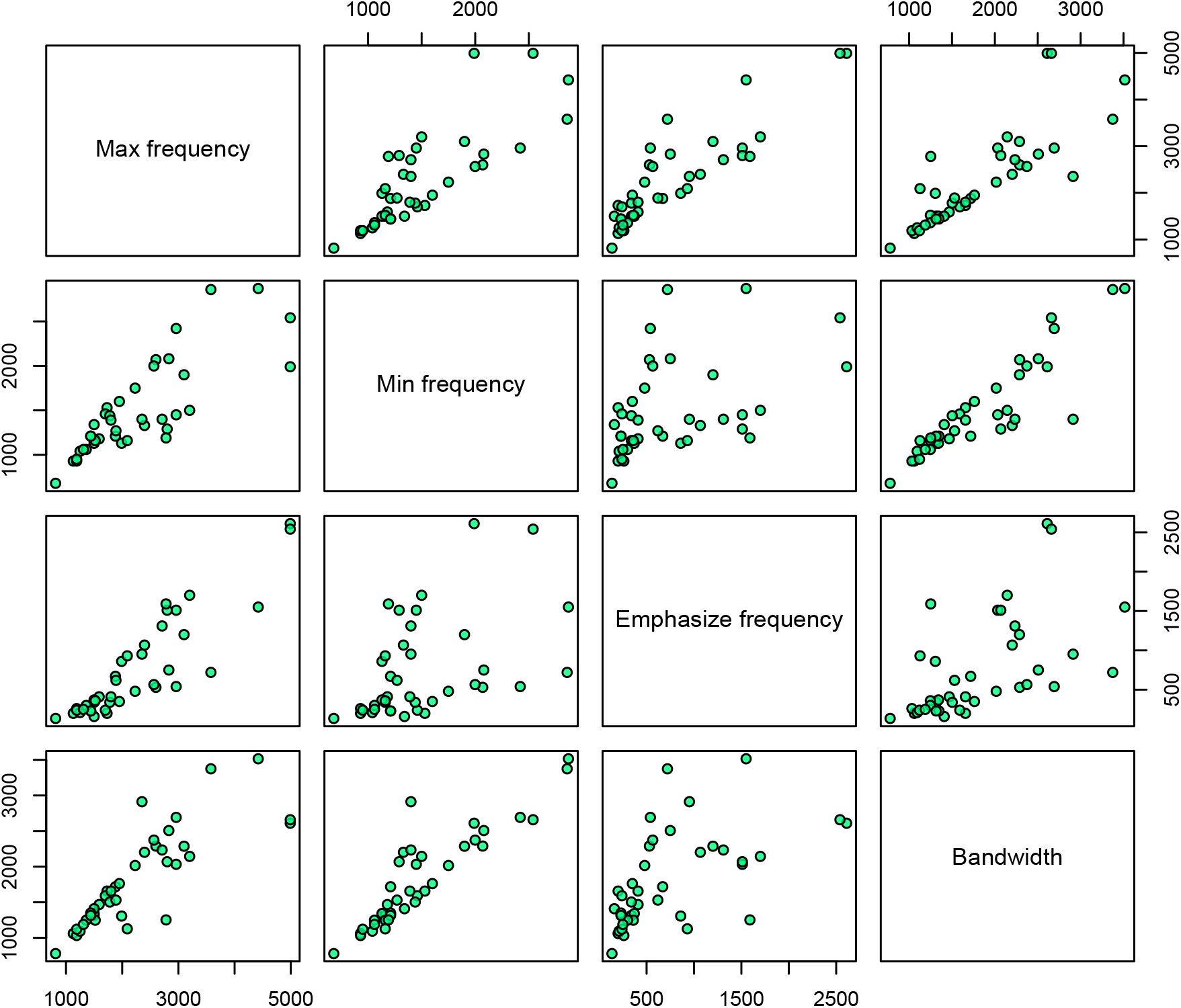
Scatterplots showing correlations among the four song variables.

